# Calcitonin gene-related peptide and intermedin induce phosphorylation of p44/42 MAPK in primary human lymphatic endothelial cells *in vitro*

**DOI:** 10.1101/2024.05.11.593644

**Authors:** Shirin R. Hasan, Dimitrios Manolis, Ewan Stephenson, Oktawia A. Ryskiewicz-Sokalska, Anthony Maraveyas, Leonid L. Nikitenko

**Affiliations:** Centre for Biomedicine, Hull York Medical School, University of Hull, Hull, UK; Hull University Teaching Hospitals NHS Teaching Trust, Queens Centre for Oncology and Haematology, Castle Hill Hospital, Hull, UK

**Keywords:** Lymphatic endothelial cells, CGRP, adrenomedullin 2, intermedin, adrenomedullin, p44/42 mitogen-activated kinases, skin

## Abstract

**Background:** Calcitonin gene-related peptide (CGRP) is a 37-amino acid peptide that belongs to the calcitonin family of peptides together with adrenomedullin (AM), adrenomedullin 2 (AM2), also known as intermedin (IMD), amylin (AMY) and calcitonin. Recent reports demonstrated the efficacy of drugs targeting CGRP signalling axis for the treatment of migraine, but also revealed significant side effects such as inflammation and microvascular complications, including aberrant neovascularisation in the skin. These findings highlight the importance of studying cellular targets of CGRP in human tissues. Recent studies using animal models implicate the role of CGRP in lymphangiogenesis and lymphatic vessel function. However, whether CGRP has direct effects on lymphatic endothelial cells (LEC) remains unknown.

**Aim:** In this study, we analysed phosphorylation of p44/42 mitogen-activated protein kinases (MAPK) in proliferating primary human dermal LEC (HDLEC) stimulated with CGRP and compared with responses to AM2/IMD and AM.

**Materials and Methods:** Primary breast HDLEC from two female donors (39 and 46 years old) were cultured *in vitro*. Cell viability and proliferation of sub-confluent HDLEC were confirmed by MTS assay and Ki67 immunostaining. Phosphorylation of p44/42 MAPK in HDLEC treated with CGRP, AM2/IMD or AM at different time points (1-30 min) and concentrations (10^-12^-10^-6^M) was assessed by immunoblotting. All experiments have been performed at least 4 times. Statistical analysis was done using Shapiro-Wilk and Kruskal Wallis tests followed by uncorrected Dunn’s test, with p<0.05 interpreted as significant.

**Results:** CGRP induced p44/42 MAPK phosphorylation in a time- and dose-dependent manner in proliferating primary HDLEC from both donors. p44/42 MAPK phosphorylation was induced by all three peptides (10^-6^M) at 5-15 min (p<0.05), but only CGRP-stimulated effect was sustained at 30 min. CGRP consistently induced p44/42 MAPK phosphorylation at 10^-7^-10^-6^M (p<0.05) in HDLEC from both donors, whilst responses to AM2/IMD and AM varied.

**Conclusions:** Our study demonstrates that CGRP and AM2/IMD have direct effects on proliferating primary HDLEC. These new findings reveal CGRP and AM2/IMD as novel regulators of lymphatic endothelial cell biology and warrant further investigation of their roles in pathologies associated with lymphatic function in the skin and other organs.

## Background/Introduction

Calcitonin gene-related peptide (CGRP) is a 37-amino acid peptide encoded by *CALCA* gene and belonging to the calcitonin family of peptides together with adrenomedullin (AM), adrenomedullin 2 (AM2), also known as intermedin (IMD), amylin (AMY) and calcitonin (Poyner et al., 2002). CGRP is a neuropeptide and a potent vasodilator that plays multiple roles in physiological and pathophysiological conditions and processes including cardiovascular disease, wound healing, inflammation, cancer, lymphoedema and migraine (Edvinsson et al., 2018; Raddant and Russo, 2011; Russo and Dickerson 2006; Recober and Russo, 2009; Russel et al., 2014; Li et al., 2013; Chai et al., 2006; Ramos-Romero et al., 2016). Recent reports demonstrated the efficacy of drugs targeting the CGRP signalling axis for the treatment of migraine patients, but also revealed significant side effects such as inflammation and microvascular complications, including aberrant neovascularisation in the skin (Wurthmann et al., 2020; Ray et al., 2021; Breen et al., 2021), highlighting the importance of studying cellular targets of CGRP in human tissues.

Recently, several studies demonstrated that CGRP plays a role in lymphangiogenesis *in vivo* (Matsui et al., 2018; Zhu et al., 2022). In *Ca/ca* knockout mice (*Ca/ca^-/-^*), lymphatic capillary formation and macrophage numbers in the skin were reduced in postoperative lymphoedematous tail model (Matsui et al., 2018). It was proposed in this study that CGRP promotes accumulation of macrophages, which produce vascular endothelial growth factor-C (VEGF-C) that acts on vascular endothelial growth factor receptor-3 (VEGFR-3) in lymphatics, leading to lymphangiogenesis (Matsui et al., 2018). In addition, in sutured mouse cornea model, exogenous CGRP induced neovascularisation and formation of larger blood and lymphatic vessels (Zhu et al., 2022). Whilst these studies demonstrated that CGRP plays a role in lymphangiogenesis, whether this peptide has direct effect on lymphatic endothelial cells (LEC) remains unknown.

The mitogen-activated protein kinases (MAPK) p44/42 are essential for angiogenesis and lymphaniogenesis (Guo et al., 2020; Bui and Hong, 2020). Previous studies demonstrated that CGRP, AM2/IMD and AM induce p44/42 MAPK phosphorylation in endothelial cells (EC) from blood vessels and endothelial progenitor cells (EPC) *in vitro* (Kim et al., 2003; Fritz-Six et al., 2008; Smith et al., 2009; Wu et al., 2018; Clark et al., 2021). In contrast to AM (Jin et al., 2008; Fritz-Six et al., 2008; Maybin et al., 2011), no studies have reported the effects of CGRP and AM2/IMD on LEC to date. To address this gap in knowledge, in the present study we investigated p44/42 MAPK phosphorylation in proliferating primary HDLEC (from two donors) cultured *in vitro* and stimulated with CGRP or AM2/IMD at different time points and concentrations, and compared and contrasted responses to AM. We found that both CGRP and AM2/IMD induce p44/42 MAPK phosphorylation in a time- and dose-dependent manner, indicating that these peptides have direct effects on LEC, and thus revealing their roles as novel regulators of LEC biology.

## Results

### HDLEC from two donors are viable and proliferative

First, we characterised HDLEC from two female donors (D1 and D2) by using lymphatic-specific and pan-EC markers (**Figure 1A**, **Supp.** Fig. 1A, B). The immunofluorescence (IF) analysis showed that HDLEC from D1 and D2 were positive for the lymphatic-specific prospero homeobox protein 1 (PROX1) and the pan-EC cluster of differentiation 31 (CD31) markers (**Figure 1A, Supp. Fig. 1A, B**). Next, the 3-(4, 5-dimethylthiazol-2-yl)-5-(3-carboxymethoxyphenyl)-2-(4-sulfophenyl)-2H-tetrazolium (MTS) assay results revealed that *in vitro* cultured HDLEC D1 and D2 were viable (**Figure 1B**). Furthermore, IF analysis demonstrated that the proliferation marker Ki67 was expressed in HDLEC from both donors (**Figure 1C**, **1D**). In summary, these findings indicate that HDLEC from D1 and D2 were positive for PROX1 and CD31, viable and proliferating *in vitro*.

**Figure 1.**
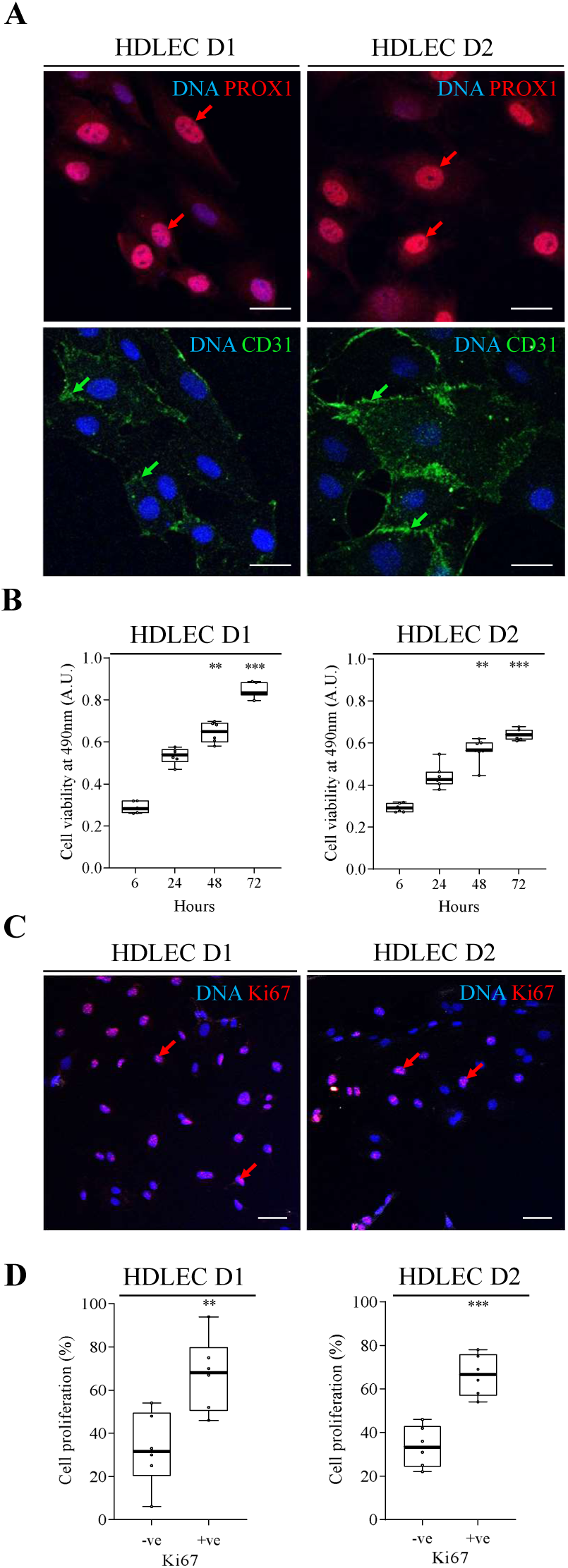
Expression of lymphatic- and pan-EC markers, cell viability and proliferation of primary HDLEC. **(A)** Characterisation of primary human dermal lymphatic endothelial cells (HDLEC) from breast skin, from two donors - D1 and D2 (see Materials & methods) was done by immunofluorescence (IF) using lymphatic- and pan-EC markers, prospero homeobox protein 1 (PROX1) (*red colour*) and cluster of differentiation 31 (CD31) (*green colour*) respectively. Nuclei were counterstained with DAPI (*blue colour*). PROX1 expression in the nuclei is shown using red arrows and CD31 expression at cell-cell contacts using green arrows. Scale bars represent 20µm. Full datasets can be found in the Supplementary Data (**Figure S1**). **(B)** Cell viability of HDLEC D1 and D2 was assessed at 6, 24, 48 and 72 hours after seeding without changing culture media. **(C)** Proliferation of HDLEC D1 and D2 were assessed by IF using mouse monoclonal antibody raised against proliferation marker Ki67 and positive staining for the marker was indicated with red arrows. Nuclei were counterstained with Hoechst (*blue colour*). Scale bars represent 50µm. Full datasets can be found in the Supplementary Data (**Figure S2**). (**B, D**) Box and whiskers plots overlaid with dots represent the results of quantification analysis of relative cell viability and proliferation. The data represents median values (n=3 independent experiments for both D1 and D2), the box contains the 25th and 75th percentiles and whiskers are the minimum and maximum values of each dataset. The statistical analysis was performed using Kruskal Wallis test (based on Shapiro-Wilk normality test) followed by uncorrected Dunn’s comparison test. *p<0.05, **p<0.01, ***p<0.001).

### CGRP and AM2/IMD induced p44/42 MAPK phosphorylation in HDLEC in a time-dependent manner

Next, we investigated whether stimulation of HDLEC D1 and D2 with CGRP, AM2/IMD and AM (all at 10^-6^M) at different time points (0-30 min) could induce p44/42 MAPK phosphorylation (**Figure 2A** and **B**). In HDLEC D1 an increase in p44/42 MAPK phosphorylation was observed after 5 minutes of exposure to each of three peptides (**Figure 2A**), compared to control (vehicle-treated cells) (**Supp.** Fig. 3). CGRP treatment of HDLEC D1 induced p44/42 MAPK phosphorylation in a time-dependent manner, starting as early as 5 minutes and with significant phosphorylation observed at 10-30 minutes (p<0.05) (**Figure 2A**). Also, significant p44/42 MAPK phosphorylation at 5-15 min (p<0.05) was observed upon AM2/IMD and AM stimulation. Overall, results for CGRP treatment were comparable to AM2/IMD and AM, but only CGRP-induced effects sustained at 30 min (**Figure 2A**). Furthermore, CGRP, AM2/IMD and AM also induced p44/42 MAPK phosphorylation in HDLEC D2 and these effects were similar to HDLEC D1 (**Figure 2B**). p44/42 MAPK phosphorylation was induced by all three peptides (10^-6^M) at 5-15 min (p<0.05), but only CGRP-stimulated effect was sustained at 30 min. In summary, our results revealed that CGRP and AM2/IMD, similar to AM, induce p44/42 MAPK phosphorylation in a time-dependent manner in HDLEC from both donors.

**Figure 2.**
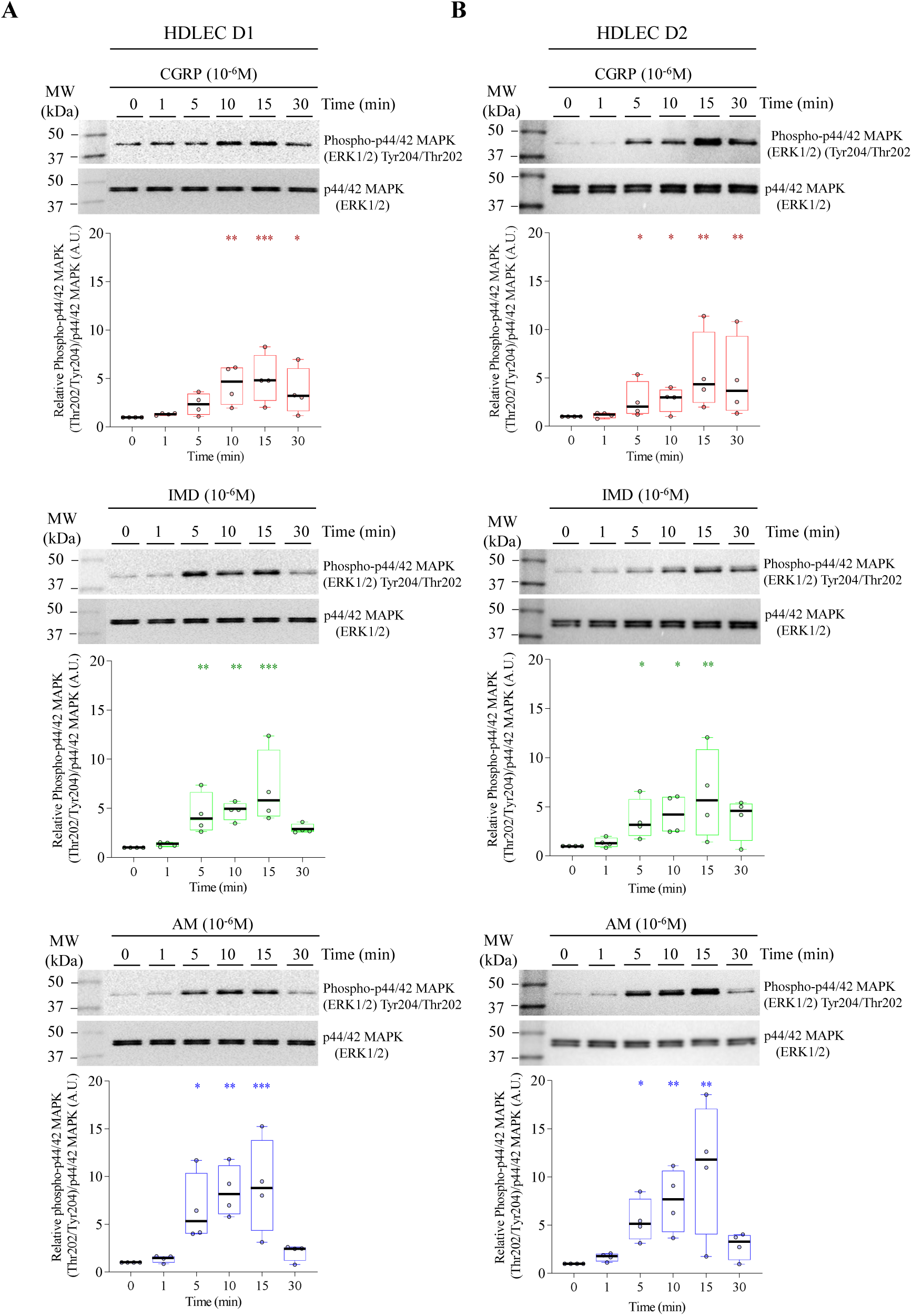
Phosphorylation of p44/42 MAPK in primary HDLEC upon stimulation with CGRP, AM2/IMD or AM at different time points. **(A, B) Dynamics of p44/42 MAPK** phosphorylation upon stimulation of primary human dermal lymphatic endothelial cells (HDLEC) from breast skin, from two donors - D1 and D2 (see Materials & methods) with calcitonin gene-related peptide (CGRP), adrenomedullin2/intermedin (AM2/IMD) or adrenomedullin (AM). Prior to cell lysis for protein extraction, HDLEC were stimulated at different time points (0-30 minutes) at 10^-6^M concentration with peptides. Phosphate-buffered saline (PBS) was used as a vehicle and a control (Supplementary Data, **Figure S3**). Bands of interest were quantified by densitometry analysis using Bio-Rad Image Lab 6.0 software. The phospho-p44/42 MAPK expression was first normalised to total p44/42 MAPK, and then to time point ’0’ for each peptide, and the results were plotted under each blot. Box and whiskers plots overlaid with dots represent the results of the quantification analysis of phospho-p44/42 MAPK relative to total p44/42 MAPK. CGRP (*red*), IMD (*green*) and AM (*blue*). The data represents median values (n=4 independent experiments for both D1 and D2), the box contains the 25^th^ and 75^th^ percentiles and whiskers are the minimum and maximum values of each dataset. The statistical analysis was performed using Kruskal Wallis (based on Shapiro-Wilk normality test) followed by uncorrected Dunn’s comparison test to determine differences compared to time point “0”. *p<0.05, **p<0.01, ***p<0.001.

### CGRP and AM2/IMD induced p44/42 MAPK phosphorylation in HDLEC in a dose-dependent manner

CGRP, AM2/IMD and AM at 10^-6^M significantly induced p44/42 MAPK phosphorylation at 10 min in HDLEC D1 and D2 (**Figure 2A, B**). Therefore, next we investigated p44/42MAPK phosphorylation at this time point in HDLEC stimulated with CGRP, AM2/IMD or AM at various concentrations (10^-12^-10^-6^M) (**Figure 3A** and **B**). The immunoblotting analysis revealed that CGRP consistently induced p44/42 MAPK phosphorylation at 10^-7^-10^-6^M in HDLEC D1 (p<0.05) and D2 (p<0.05) (**Figure 3A** and **B**). These effects of CGRP were comparable to AM2/IMD (10^-7^-10^-6^M) and AM (10^-7^-10^-6^M)-stimulated HDLEC D1 (**Figure 3A**). In HDLEC D2, AM2/IMD and AM induced p44/42 MAPK phosphorylation at 10^-8^-10^-6^M and 10^-9^-10^-6^M, respectively (**Figure 3B**). CGRP consistently induced p44/42 MAPK phosphorylation at 10^-7^-10^-6^M (p<0.05) in HDLEC from both donors, whilst responses to AM2/IMD and AM varied. In summary, our results revealed that CGRP and AM2/IMD, similar to AM, induce p44/42 MAPK phosphorylation in a dose-dependent manner.

**Figure 3.**
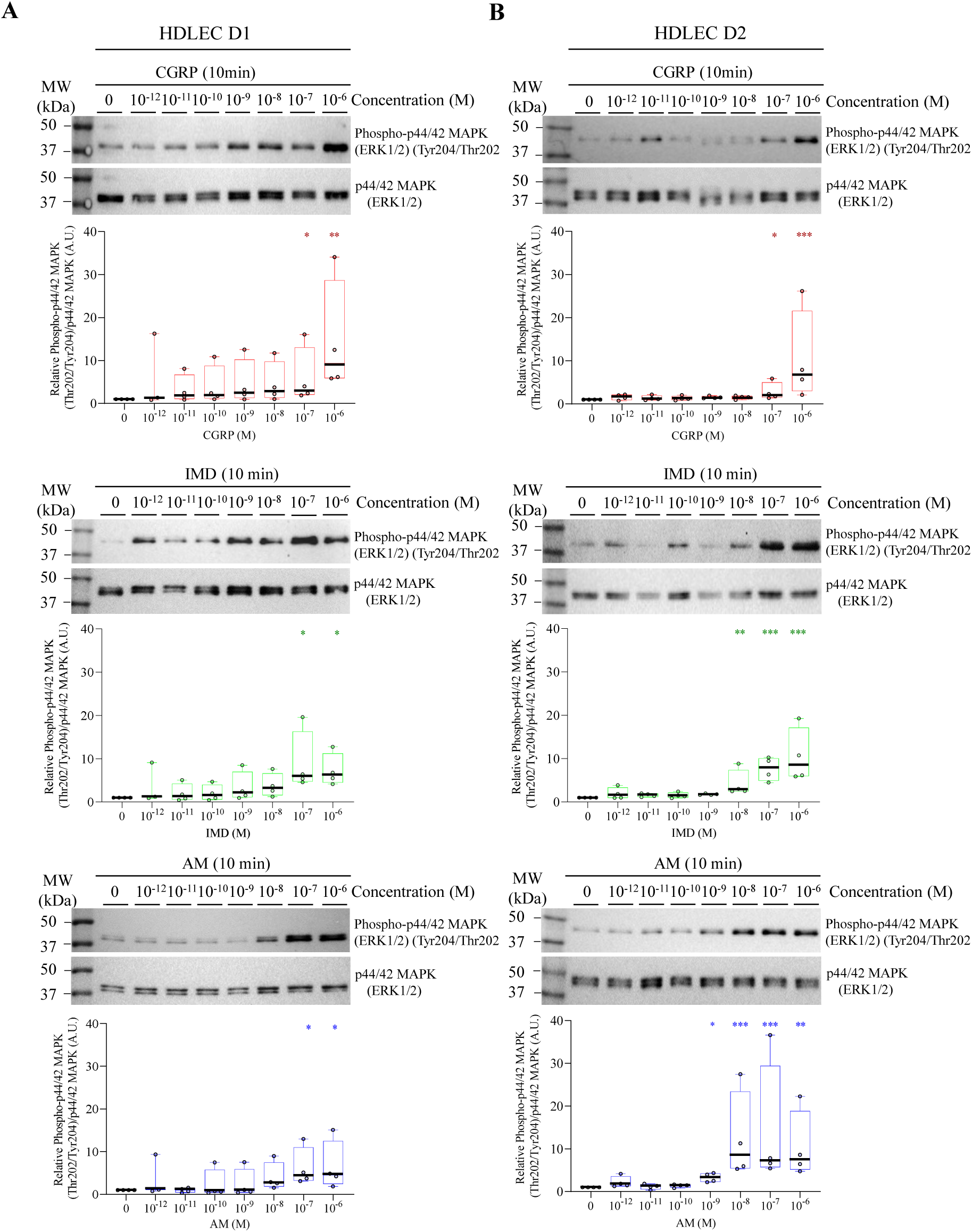
Phosphorylation of p44/42 MAPK in primary HDLEC upon stimulation with CGRP, AM2/IMD or AM at different concentrations. **(A, B)** Dynamics of p44/42 MAPK phosphorylation upon stimulation of primary human dermal lymphatic endothelial cells (HDLEC) from breast skin, from two donors - D1 and D2 (see Materials & methods) with calcitonin gene-related peptide (CGRP), adrenomedullin2/intermedin (AM2/IMD) or adrenomedullin (AM). Prior to cell lysis for protein extraction, HDLEC were stimulated with peptides at various concentrations (10^-12^-10^-6^M) for 10 minutes. Phosphate-buffered saline (PBS) was used as a vehicle. Bands of interest were quantified by densitometry analysis using Bio-Rad Image Lab 6.0 software. The phospho-p44/42 MAPK expression was first normalised to total p44/42 MAPK, and then to vehicle (PBS) for each peptide, and results were plotted under each blot. Box and whiskers plots overlaid with dots represent the results of the quantification analysis of phospho-p44/42 MAPK relative to total p44/42 MAPK. CGRP (*red*), IMD (*green*) and AM (*blue*). The data represents median values (n=4 independent experiments for both D1 and D2), the box contains the 25^th^ and 75^th^ percentiles and whiskers are the minimum and maximum values of each dataset. The statistical analysis was performed using Kruskal Wallis (based on Shapiro-Wilk normality test) followed by uncorrected Dunn’s comparison test to determine differences between stimulated and unstimulated cells. *p<0.05, **p<0.01, ***p<0.001.

## Discussion

CGRP is involved in a variety of biological processes such as cardiovascular homeostasis (Favoniyq et al., 2019), immune regulation (Wallrapp et al., 2019), pain modulation (Iyengar et al., 2017) and, most recently, lymphangiogenesis (Matsui et al., 2018; Zhu et al., 2022). In the present study, we found that CGRP induced p44/42 MAPK phosphorylation in a time- and dose-dependent manner in proliferating primary HDLEC from two donors. To our knowledge, this is the first of its kind report about the direct effect of CGRP on LEC biology. This novel finding warrants further investigation of CGRP role in these cells in the context of conditions and pathologies associated with lymphatic function and therapies targeting CGRP signalling axis.

In animal models, CGRP is proangiogenic during hind limb ischaemia (Zheng et al., 2010), tumour growth (Toda et al., 2008) and wound healing (Toda et al., 2008b), and prolymphangiogenic in the sutured cornea (Zhu et al., 2022) and postoperative tail models (Matsui et al., 2018). In the present study, we found that CGRP and AM2/IMD have direct effects on proliferating primary HDLEC (from two donors) cultured *in vitro*. Besides embryogenesis, LEC proliferation also takes place in adulthood in some physiological processes (wound healing and inflammation) and pathological conditions (cancer and lymphoedema) (Toda et al., 2008b; Toriyama et al., 2015; Toda et al., 2008; Matsui et al., 2018; Zhu et al., 2022). Under these processes and conditions, the expression and role of CGRP has been also reported based on findings from *in vivo* models (Toda et al., 2008; Toda et al., 2008b; Matsui et al., 2018; Zhu et al., 2022). For example, in *Ca/ca^-/-^* knockout mice microvessels density and area, and M2 macrophage numbers were reduced during wound healing in the skin, lung cancer and postoperative lymphoedema (Toda et al., 2008; Toda et al., 2008b; Matsui et al., 2018). In addition, in a laser-induced choroidal neovascularization model, inflammation was suppressed in *Ca/ca^-/-^* knockout mice (Toriyama et al., 2015). Several studies established the association of inflammation with lymphangiogenesis, lymphatic vessel remodelling and function (Baluk et al., 2009; Schwager and Detmar, 2019). Therefore, since CGRP has been implicated in neovascularisation and action on microvasculature under conditions where lymphangiogenesis takes place, the findings from our *in vitro* study suggest that CGRP might have direct effects on proliferating LEC *in vivo* too.

Phosphorylation of p44/42 MAPK in EC is crucial during angiogenesis and lymphangiogenesis (Kim et al., 2003; Bui and Hong, 2020). Previous studies established a link between the MAPK signalling pathway and members of the calcitonin family of peptides, including CGRP, in several cell types, such as human umbilical vascular endothelial cells (HUVEC) and EPC, cardiomyocytes (Clark et al., 2021; Wu et al., 2018), Schwann cells (Permpoonputtana et al., 2016) and smooth muscle cells (Schaeffer et al., 2003). In the present study, we found that CGRP and AM2/IMD, comparably to AM, induce p44/42 MAPK phosphorylation in proliferating primary HDLEC cultured *in vitro*. Our findings revealed that CGRP consistently induced phosphorylation of p44/42 MAPK at 10^-7^-10^-6^M in HDLEC obtained from breast skin from two adult female donors. These results are in agreement with other reports, which demonstrated that p44/42 MAPK phosphorylation was induced by CGRP at 10^-7^-10^-5^M in EC of embryonic origin (HUVEC and EPC) (Clark et al., 2021; Wu et al., 2018). Furthermore, CGRP induces other signalling pathways, including protein kinase B (Akt) in human dermal microvascular EC (Nikitenko et al., 2006), endothelial nitric oxide (NO) in keratinocytes (Yapping et al., 2006), cAMP, NO and calcium (Ca^2+^) mobilisation in HUVEC (Clark et al., 2021). Therefore, CGRP and AM2/IMD effects on various signalling pathways in proliferating primary human LEC from adult tissues (both male and female donors) require further investigations.

Proliferating LEC acquire quiescence once the lymphatic network is established (Petrova et al., 2020; Oliver et al., 2020). In the quiescent state, LEC are involved in immune cell transportation and other functions of lymphatic vessels (Petrova et al., 2020; Oliver et al., 2020). Findings form several studies suggest that CGRP plays a role in quiescent lymphatic vessels (Yamada and Hoshino, 1996; Hosaka et al., 2006; Ding et al., 2016; Wang et al., 2018). For example, CGRP-containing nerve fibres are located in close proximity to lymphatic capillaries in rat skin (Yamada and Hoshino, 1996). In addition, exogenous CGRP stimulation leads to inhibition of NO-dependent vasomotion in perfused mesenteric lymphatic vessels of guinea pigs (Hosaka et al., 2006). Also, *in vitro* studies demonstrated that by acting on primary murine dermal microvascular EC, CGRP facilitates the immune response and regulates the outcome of antigen presentation by Langerhans cells to T cells (Ding et al., 2016). Furthermore, studies using *Adm2^-/-^*knockout mice, implicated AM2/IMD in the proliferation of quiescent CD31-positive vessels in the retina (Wang et al., 2018). Therefore, it cannot be excluded that observed in these studies effects of CGRP and AM2/IMD may be due to direct action of these peptides on quiescent LEC in lymphatic vessels, thus warranting further investigation.

CGRP effects on cells are mediated by G-protein coupled receptors (GPCRs) calcitonin receptor-like receptor (CLR) or calcitonin receptor (CTR) upon their co-expression with receptor activity-modifying proteins 1, 2 and 3 (RAMPs 1-3) (Piozak and Hay, 2020). CLR and RAMPs 1-3 expression was previously shown in primary human LEC in tissues (skin, uterus and lung) and *in vitro* (Vart et al., 2007; Jin et al., 2008; Maybin et al., 2011; Nikitenko et al., 2013; Faulkner et al., 2023; Manolis et al., 2023). CGRP and AM compete for the same receptor in human dermal microvascular EC, involving different downstream mechanisms (Nikitenko et al., 2006). Furthermore, our recent study revealed that endogenously expressed in HDLEC CLR interacts with 37 novel proteins, which are involved in signalling, post-translational modifications, and trafficking of other GPCRs (Manolis et al., 2023). Altogether, these findings suggest that further studies are needed to dissect the role of CLR (or CTR) in mediating CGRP- and AM2/IMD-induced direct effects in primary LEC that were revealed in our study.

Recent reports demonstrated the use of drugs targeting the CGRP signalling axis, such as antibodies against CGRP and its receptors, for the effective treatment of migraine (Holland et al., 2018; Edvinsson et al., 2018; Majima et al., 2019; Labastida-Ramirez et al., 2023). Since CGRP is implicated in both lymphangiogenesis and angiogenesis (Toda et al., 2008; Zheng et al., 2010; Matsui et al., 2018; Zhu et al., 2022), it was proposed that prolonged treatment with these agents could lead to impaired neovascularisation in some patients (Majima et al., 2019). Recent studies demonstrated that the use of CGRP monoclonal antibody (erenumab) resulted in skin disturbance during wound healing in a migraine patient, with skin biopsy demonstrating a deep perivascular and interstitial lymphohistiocytic infiltrate with admixed eosinophils, ulceration of the epithelium, heavy oedema of the papillary dermis and focally thrombosed vessels (Wurthmann et al., 2020). In addition, eight migraine patients treated with CGRP monoclonal antibodies (erenumab, fremanezumab or galcanezumab) developed inflammatory complications (Ray et al., 2021; Gross et al., 2019). These complications include Susac’s syndrome (linked to endotheliopathy, a disorder that develops due to functional changes in the endothelium) (Gross et al., 2019), granulomatosis with polyangiitis (a rare disorder that causes inflammation of the blood vessels in the nose, sinuses, throat, lungs and kidneys), drug reaction with eosinophilia and systemic symptoms (clinically presents, as an extensive mucocutaneous rash, accompanied by fever, lymphadenopathy, hepatitis, haematologic abnormalities with eosinophilia and atypical lymphocytes), autoimmune-hepatitis, poly-arthralgia, psoriasis and urticarial eczema (Ray et al., 2021). All these effects were *de novo*, with a clear temporal relationship between exposure and symptom-onset (Ray et al., 2021). Moreover, in a retrospective cohort study of 169 migraine patients treated with CGRP monoclonal antibodies (erenumab, fremanezumab or galcanezumab), nine patients exhibited microvascular complications (Breen et al., 2021). In the context of these reports, the findings from our study suggest that adverse/side effects of drugs targeting the CGRP signalling axis could be attributed (at least partially) to CGRP signalling and functions in human LEC and lymphatic vessels.

In summary, our first of its kind study reveals that CGRP and AM2/IMD induce phosphorylation of p44/42 MAPK in proliferating primary HDLEC, and thus have direct effects on these cells. These new findings suggest that CGRP and AM2/IMD are novel regulators of LEC biology. Our study opens up new avenues and serves as a springboard for further investigations of CGRP- and AM2/IMD-induced effects in both proliferating and quiescent LEC (and also other cellular targets for these peptides in human tissues) under physiological conditions and pathologies associated with lymphatic function and beyond.

## Materials and Methods

### Reagents

Synthetic human adrenomedullin (AM) trifluoroacetate salt (#4030315), human a-calcitonin gene-related peptide (CGRP) (#4013281) and human intermedin (IMD) (#4044529) were from Bachem. Primary and secondary antibodies were obtained from a range of manufacturers and used at dilutions and concentrations described below. Rabbit polyclonal anti-human CLR (LN-1436; 1:1000) was raised and characterised previously (Nikitenko et al., 2006). Primary mouse monoclonal CD31/platelet adhesion molecule-1 (PECAM-1 (#555444; 1:100), mouse IgG1 (#555746; 1:100) and mouse IgG2 (#555740; 1:100) all from BD Biosciences. Mouse monoclonal beta-actin (#ab6276; 1:1000) was from Abcam, rabbit polyclonal anti-phosphorylated p44/42 MAPK (#9101S; 1:1000) and anti-total 44/42 MAPK (#9102S; 1:1000) were from Cell Signalling Technology, goat polyclonal PROX1 (#AF2727; 1:100), rabbit IgG (#ab105C; 1:1000) and goat IgG (#ab108C; 1:200) were from R&D Systems. Secondary conjugated polyclonal donkey anti-rabbit IgG Alexa 594 (#A-21206; 1:600), anti-rabbit IgG Alexa 488 (#A-21207; 1:600), anti-mouse IgG Alexa 488 (#A-21202; 1:600), anti-goat IgG Alexa 488 (#A-11055; 1:600), anti-goat IgG Alexa 594 (#A-11058; 1:600) and phalloidin Alexa Fluor™ 635 (#A34054; 1:100) all from Invitrogen, goat horseradish peroxidase (HRP) anti-mouse IgG (#P0447; 1:1000) and anti-rabbit IgG (#P0448; 1:1000) were from Dako.

### Ethical statement and cell culture

#### Ethica/statement

Primary human dermal lymphatic endothelial cells (HDLEC, passage 1) from breast skin of two healthy female donors (46-year-old-D1 and 39-year-old-D2) were purchased commercially from PromoCell® (lot #431Z012.3 and #406Z043.4 respectively). These cells have been tested by the manufacturer for the absence of HIV-1, HIV-2, HBV, HCV, HTLV-1, HTLV-2 and microbial contaminants and by us for the absence of mycoplasma using EZ-PCR Mycoplasma Test kit (Biological Industries; #20-700-20) and the HyperLadder^TM^ 1kb (Bioline/Meridian Bioscience; # BIO-33026).

#### Cell culture

Throughout this study HDLEC from D1 and D2 were used in sub-confluent (proliferating) conditions. Cells were seeded onto T-75 (Greiner Bio One; #658950) pre-coated flask in microvascular (MV2) EC full growth medium (PromoCell®; #C-22121). Recombinant human vascular endothelial growth factor-C (VEGF-C) (R&D Systems; #9199-VC; 7.5 ng/mL) was also added to the MV2 full growth medium. Cultures were incubated at 37°C in a 5% CO_2_ humidified atmosphere, and the medium was replaced every 24 hours. Cells were passaged 1:3 at confluence (~80%) by release with trypsin/EDTA (ethylenediaminetetraacetic acid) and sub-cultured using the same method.

### Immunofluorescence

HDLEC characterisation was done by using immunofluorescence as previously described (Nikitenko et al., 2006). HDLEC from D1 and D2 were seeded (5000 cells per well) into 8-well slide chambers (Fisher Scientific; #16250681). Once cells reached 80% confluency, HDLEC were starved in MV2 basal growth medium containing 0.5% foetal bovine serum (FBS (Gibco; #10500064). After 24 hours, cells were fixed using either 4% paraformaldehyde (PFA) or acetone/methanol (2:3 ratio) solutions. Prior to 4% PFA fixation for seven minutes, the culture medium was removed, and cells were gently washed with phosphate-buffered saline (PBS) (Fisher Scientific; #10209252). Next, the PFA solution was removed and cells were washed with PBS once and stored in PBS at 4°C until further experimentation. For fixation using acetone/methanol, the culture medium was removed, and cells were gently washed with PBS once and incubated with the fixative for three minutes. Next, the fixative was removed and the plates were left to air dry for 20-30 minutes, then slides were wrapped in cling film and stored at -20°C. Prior to primary antibody incubation, a pre-blocking step, using 10% donkey serum (Bio-Rad; #C06SB, diluted in PBS/Triton-X100) for 30 minutes at room temperature (RT) was performed. Primary antibodies were diluted at appropriate concentrations in 2% donkey serum and added to 8-well slides, wrapped in cling film and incubated overnight at 4°C. The next day, cells were placed on ice and washed with PBS/Triton three times for three minutes before the incubation with secondary antibodies in 2% donkey serum solutions. Incubation with secondary fluorophore-conjugated antibodies was performed under light protection at RT for 45 minutes. Next, the secondary antibody solution was removed and cells were washed with PBS/Triton-X100 thrice. When required, incubation with phalloidin (Alexa FluorTM 635) was performed for 40 minutes at RT, followed by mounting using an antifade mounting medium 4’,6-diamidino-2-phenylindole (DAPI, Vectashild®, Vibrance; #H15800-2). Cells were imaged using the confocal microscope (ZEISS LSM 710) and x20 objective. Image analysis was performed using the Zen Blue 3.0 (ZEISS) software.

### Cell viability assay

HDLEC form D1 and D2 were sub-seeded (3500 cells) into 96-well plates (SARSTEDT; #83.3920) in MV2 full growth medium. 24 hours post-sub-seeding, HDLEC from D1 and D2 were starved in MV2 basal growth medium containing 0.5% FBS. After 24 hours a 3-(4, 5-dimethylthiazol-2-yl)-5-(3-carboxymethoxyphenyl)-2-(4-sulfophenyl)-2H-tetrazolium (MTS) assay was performed according to the manufacturers protocol. Briefly, 20µL of CellTiter-96®Aqueous and One Solution MTS assay (Promega; #G3580) was added to each well using a multichannel pipette (Transferpette, SLS Labs) and plates were incubated for 120 minutes at 37°C and 5% CO_2_. The measurements of absorbance at a wavelength of 490.0 nm were taken using a Tecan Infinite M200 Plate Reader.

### Stimulation with peptides

Peptide stimulation of HDLEC from D1 and D2 was performed as previously described (Nikitenko et al., 2006). The cells were sub-seeded (~75000 cells per well) into 6-well plates (SARSTEDT; #83.3924). The next day, cells were starved in MV2 basal growth medium containing 0.5 % FBS. After 24 hours HDLEC form Dl and D2 were treated with the same volume (15µL) of peptides (AM, AM2/IMD or aCGRP at 10^-6^ M, in filtered PBS or vehicle (PBS) at different time points (0-30 minutes) and at different concentrations (10^-12^-10^-6^M) for 10 minutes. Next, 6-well plates were placed at 37°C in a 5% CO_2_ humidified atmosphere for the indicated time point and then processed for cell lysis.

### Cell lysis

HDLEC (Dl and D2) lysis was performed as previously described (Nikitenko et al., 2006). All steps were performed on ice. Cells were washed with ice-cold filtered PBS and lysates were harvested using cell scrapers in radioimmunoprecipitation assay (RIPA) lysis buffer solution, in which phosphatase inhibitor (phosSTOP^TM^, Roche; #4906845001) was added. Samples were processed in 1.5 ml tubes aspirating up and down and repeating three times at 10-minute intervals. Insoluble material was pelleted by centrifugation at 13,000*g* for 10 minutes at 4°C, and the supernatant was stored at -20°C prior to the determination of total protein concentration.

### Quantification of total protein concentration

Bicinchoninic acid (BCA) assay (Pierce™; #10678484) was used for the determination of total protein concentration of cell lysates according to the manufacturer’s instructions. The measurements of absorbance at a wavelength of 562.0 nm were taken using a Tecan Infinite M200 Plate Reader.

### SDS-PAGE and immunoblotting

Protein lysates from HDLEC D1 and D2 were subjected to sodium dodecyl sulphate-polyacrylamide gel electrophoresis (SDS-PAGE) and immunoblotting as previously described (Nikitenko et al., 2006). Samples were electrophoretically separated on 10% polyacrylamide-based gel (acrylamide/methylene bisacrylamide solution at 37.5:1 ratio, 375mM Tris-pH8.8, 0.1% SDS, 0.l% ammonium persulfate (APS) and 0.04% tetramethylethylenediamine (TEMED) set with 5% stacking gel (acrylamide/methylene bisacrylamide solution at 37.5:1 ratio, 126 mM Tris-pH6.8, 0.l% SDS, 0.1% APS and 0.01% TEMED). Electrophoresis was run using Tris/Glycine/SDS-based (0.25M Tris, 1.92M Glycine, 800mL double distilled water, 1%SDS, pH8.3) running buffer at 100V for 120 minutes or until optimal resolution of proteins was obtained at 4°C. Transfer to polyvinylidene difluoride (PVDF) membrane (GE Healthcare Life Science, Amersham; #1060002l Xl) was performed using Tris/Glycine-based (0.25M Tris, 1.92M Glycine, 800mL double distilled water, pH8.3) transfer buffer at 60V for three hours at 4°C. The membranes were incubated in a blocking solution (5% non-fat milk in Tris-buffered saline containing 0.5% Tween-20 (TBS/T) for 60 minutes prior to primary antibody incubation. For primary antibody incubation, the membranes were placed in a 50mL falcon tube (SARSTEDT; #62.547.254) containing the primary antibody solution and incubated overnight on a tube roller at 4°C. For secondary HRP-conjugated antibody incubation, membranes were washed with TBS/T three times for 5 minutes and added to a tube containing the relevant secondary antibody and incubated at RT for 45 minutes. HRP activity was then detected using an enhanced chemiluminescence (ECL) kit (Bio-Rad; #170-5061). After detection, the membranes were stripped by using a stripping buffer (Fisher Scientific; #10016433) and re-probed or stored at -20°C. Anti-human B-actin was used as a control to monitor and confirm equal loading of total protein in samples. Imaging and densitometry were performed using Bio-Rad ChemiDoc XRS+ System (BioRad Laboratories, Herefordshire) and Bio-Rad ImageLab software (version 6.1) respectively. Exposure times were varied and relied on the quality and intensity of the obtained signal.

### Statistical analysis

All results were presented as box plots with median values with maximum and minimum whiskers. Immunoblotting images for p44/42 MAPK were analysed in ImageLab software using the raw images obtained from the Chemidoc^TM^ XRS + System. The relevant bands were identified/selected (p44/42MAPK and total p44/42 MAPK) by drawing a box around the appropriate area and the signal within that area was quantified by using the software (version 6.1). Phospho-p44/42 band intensity data were normalized to the intensity of the total p44/42 MAPK, then normalized throughout to the untreated “0” minute/concentration of p44/42 MAPK band. The Shapiro-Wilk normality test was used to test the normal distribution of the data set. The non-parametric Kruskal Wallis test was performed, followed by the uncorrected Dunn’s test, which compares each group to the control group, without any corrections for multiple comparisons. Results were deemed significant if p<0.05. GraphPad Prism 8 software (San Diego, CA, USA) was used for statistical analysis unless stated otherwise. The different statistical tests used for relevant experiments are described in individual figure legends.

## Acknowledgements

Mr. Eamon Faulkner and Dr. Matthew Morfitt for their constructive feedback on this manuscript. Funding: University of Hull Research Support Fund (E.S. and L.L.N.); Welcome Trust Biomedical Vacation Scholarship (Reference: 218420/Z/19/Z; D.M. and L.L.N.); Biochemical Society UK Summer Vacation Studentship (O.A.R-S and L.L.N.); and The Melanoma Fundable Charity Castle Hill Hospital (S.R.H., L.L.N. and A.M.).

## Author contributions

Conceptualisation and Design, L.L.N.; Methodology, S.R.H., and L.L.N.; Immunoblotting and Image analysis, S.R.H., D.M., E.S., O.A.R-S., and L.L.N.; Immunofluorescence and Image analysis, S.R.H., D.M., and L.L.N.; Data Analysis, Statistics and Interpretation, S.R.H. and L.L.N.; Writing, Review and Editing, S.R.H., A.M. and L.L.N.; Supervision, A.M. and L.L.N.

