## Supplementary Figure Legends for "Calcitonin gene-related peptide and intermedin induce phosphorylation of p44/42 MAPK in primary human lymphatic endothelial cells *in vitro*"

**Figure S1. Expression of lymphatic and pan-EC specific markers in primary HDLEC.** Characterisation of primary human dermal lymphatic endothelial cells (HDLEC) from breast skin from two donors-D1 and D2 (see Materials & methods) was done by immunofluorescence (IF) using lymphatic-specific prospero homeobox protein 1 (PROX1) and pan-endothelial cluster of differentiation 31 (CD31) markers, and then imaged using ZEISS LSM 710 confocal microscope (*left panel*). Goat (G) and mouse (M) immunoglobulins (IgG) were used as species-matching controls for primary antibodies (*right panel*). Conjugated donkey anti-goat (Alexa 594) and anti-mouse (Alexa 488) secondary antibodies were used. Nuclei were counterstained with DAPI (*blue colour*). Red and green arrows indicate positive staining for individual antigens. The bottom right image represents the merged channels (*left panel*), and only merged channels are represented for controls (*right panel*). Scale bars represent 20µm.

**Figure S2. Expression of Ki67 proliferation marker in primary HDLEC.** Proliferation of primary human dermal lymphatic endothelial cells (HDLEC) from donor (**A**) D1 and (**B**) D2 were analysed by immunofluorescence using mouse monoclonal antibody raised against proliferation marker Ki67 and conjugated donkey anti-rabbit (Alexa 594) secondary antibody (*red colour*) and phalloidin (binding F-actin) conjugated to Alexa 635 (*white pseudo-colour*) and imaged using ZEISS LSM 710 confocal microscope. Nuclei were counterstained with Hoechst (*blue colour*). Positive staining was indicated with red arrows. Scale bar represents 50µm.

**Figure S3. Phosphorylation p44/42 MAPK in primary HDLEC upon PBS stimulation at different time points. (A, B)** Dynamics of p44/42 MAPK phosphorylation upon stimulation of primary human dermal lymphatic endothelial cells (HDLEC) from donors D1 and D2 (see Materials & methods) with phosphate-buffered saline (PBS), used as a vehicle and control for peptide stimulation (**Figure 2**). Prior to cell lysis for protein extraction, HDLEC were stimulated with PBS at different time points (0-30 minutes). Bands of interest were quantified by densitometry analysis using Bio-Rad Image Lab 6.0 software. The phospho-p44/42 MAPK expression was first normalised to total p44/42 MAPK, and then to time point '0' and results were plotted under each blot. Box and whiskers plots overlaid with dots represent the quantification analysis of phospho-p44/42 MAPK relative to total p44/42 MAPK. The data represents median values for both (independent experiments for both D1 and D2), the box contains the 25th and 75th percentiles and whiskers are the minimum and maximum values of each dataset. The statistical analysis was performed using Kruskal Wallis (based on Shapiro-Wilk normality test) followed by uncorrected Dunn's comparison test to determine differences compared to time point "0". *p<0.05, **p<0.01, ***p<0.001.
