## Supplementary figures and images for "Calcitonin gene-related peptide and intermedin induce phosphorylation of p44/42 MAPK in primary human lymphatic endothelial cells *in vitro*"

### Supplementary Figures

**Supplementary Figure 1.**

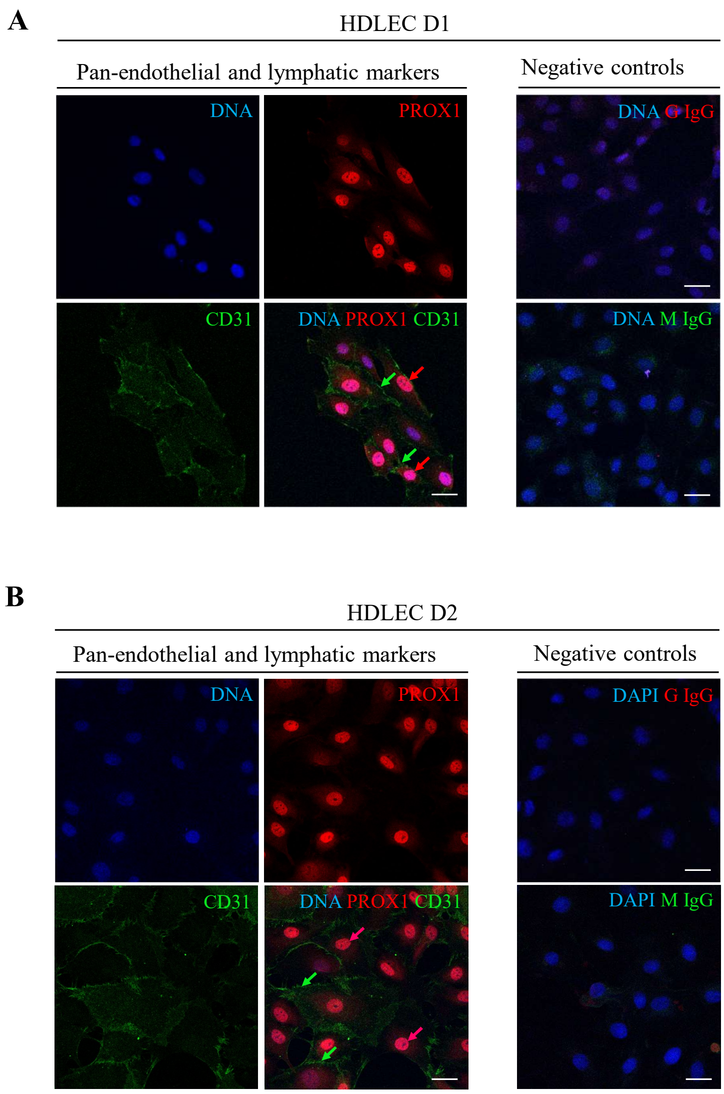

**Supplementary Figure 2.**

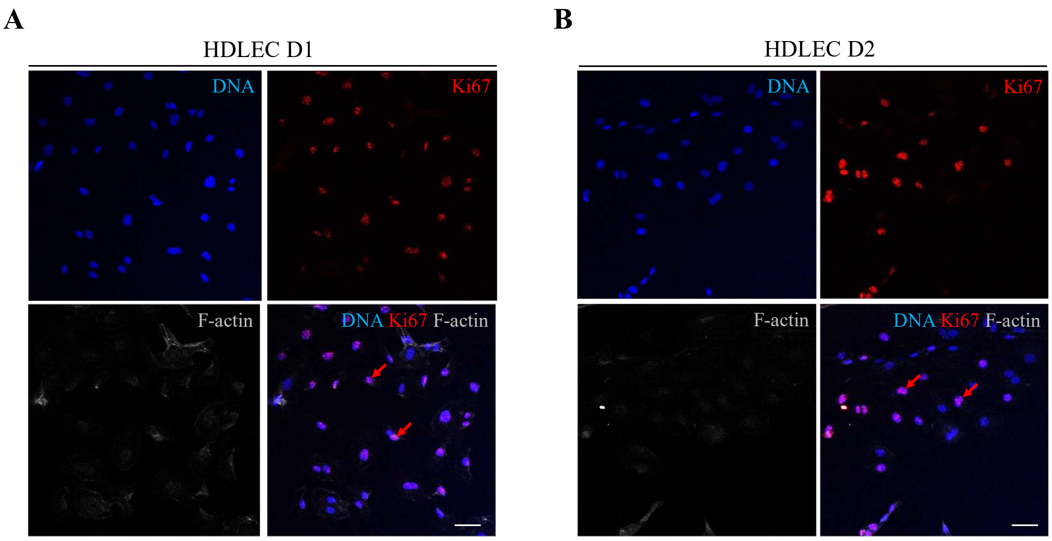

**Supplementary Figure 3.**

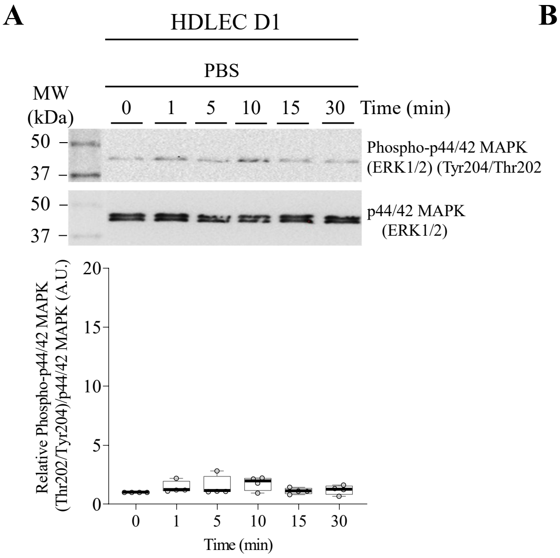


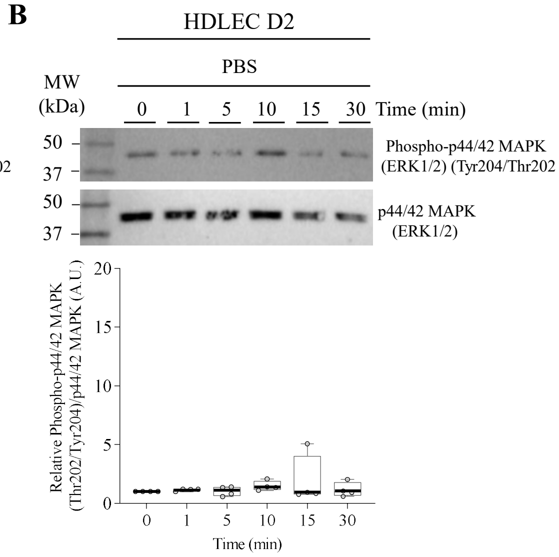
